# The concept of the gain curve

**DOI:** 10.1101/2024.01.07.574552

**Authors:** Martin Burd

## Abstract

Gain curves were introduced to explain how hermaphrodites could displace a dioecious population, and to account for sexual allocation in hermaphrodites. Terms for gamete production employed for the first purpose were transformed for the second into male and female gain curves that ostensibly defined fitness outcomes. These gain curves pose a conceptual challenge if they are specified separately because fitness at the population level cannot occur through one sex function independently of success through the other. If gain curves truly represent fitness outcomes, anomalies can arise, such as inequality of total male and female fitness in a population. Gain curves were originally used in a mathematical framework that treated the ostensible gain functions as inputs of male and female actors to a mating arena rather than as mating outcomes from that arena. I present a model of sex allocation that incorporates power functions to describe both gamete production and fitness gain in a manner that explicitly separates these two roles. In this formulation, the gamete production functions have the identical effect on optimal sex allocation originally attributed to gain curves while the true fitness gain curves lose nearly all effect on the optimum. Thus, despite the label, gain curves were implicitly describing inputs rather than outcomes. Because gain curves have been a staple of evolutionary ecology for decades, the implication is that much of our understanding of sexual allocation in hermaphrodites needs to be revisited. I outline some directions such an effort might take.

## Introduction

A gain curve represents a pattern of fitness accrual in response to resource investment in male or female function by hermaphroditic organisms. The idea of a fitness gain curve is simple and inevitable. If no resources are devoted to reproduction, no fitness can be attained. As resources investment rises, there must be some corresponding fitness—even potentially zero fitness— yielding a function on a Cartesian quadrant in which the horizontal axis represents resource investment (or production of mating entities like sperm and eggs, microspores and megaspores, or pollen and ovules) and the vertical axis represents fitness. The function may be linear or nonlinear; it might not be monotonic; but some function that includes the origin must correspond to the pattern of fitness acquisition. As a matter of description, then, gain curves are inescapable. But gain curves are thought to be more than a convenient descriptive shorthand for a feature of reproductive ecology. They are meant to explain sexual allocation in hermaphrodites.

Gain curves were introduced by Charnov (1979, 1982) and have been widely adopted as an element of sex allocation theory (Charlesworth and Charlesworth 1981; Charnov and Bull 1986; Morgan 1992; Sakai 2000; Klinkhamer and de Jong 2002; Sato 2002; Zhang and Jiang 2002; de Jong et al. 2008) and as a framework for interpreting empirical data on reproductive success in hermaphroditic plants and animals (Emms 1996; Emms et al. 1997; Campbell 2000; Baeza 2007; Vizoso and Schärer 2007; Johnson and Yund 2009; Rosas and Domínguez 2009; Perry and Dorken 2011; Vellnow et al. 2018). Despite the wide acceptance that gain curves have received, I argue here that they lack explanatory power. This suggestion will be startling after decades in which gain curves have been a foundation of the theory. I therefore propose to step through Charnov’s (1982) original derivation of his hermaphroditic sex allocation model to demonstrate why gain curves lack the effect on sex allocation that was attributed to them. Functions that ostensibly represent fitness gains were used in a way that actually represents the input of male and female gametes into a mating arena rather than the fitness outcome of their interaction. Having used gain curves in this way, it is then possible to calculate an optimal sex allocation, but the putative optimum continues to “assume” that the functions represent inputs. When they are nonetheless interpreted as fitness outcomes, anomalies can arise, such as unequal total male and female fitness at the population level. Inequality is unobjectionable for the inputs to a mating process—sperm or pollen commonly outnumber eggs or ovules—but is clearly illegitimate as an outcome.

Below I derive Charnov’s (1982) original gain curve argument to show how the conflation of inputs and fitness outcomes arose. I then use power-function gain curves in a way that truly influences the fitness outcome, a use that avoids inconsistencies inherent in the formulation of Charnov (1982), but the true gain curves have almost no effect on optimal sex allocation.

### Gamete production and fitness consequences

Charnov (1979, 1982; also Charnov et al. 1976) was concerned with the conditions that favored either hermaphroditism or dioecy as well as the division of reproductive resources between the male and female functions of hermaphrodites. His development referred to flowering plants and the production of pollen and seeds (via ovules), but the argument works equally for any male and female mating agents, either gametes or more complex entities like microspores and megaspores or pollen and ovules that themselves produce gametes. I will retain the focus on flowering plants and pollen and ovules to be concrete.

#### Production

Charnov (1982, Chapter 14) supposed that a population consists of *n*_1_ males, *n*_2_ females, and *n*_3_ hermaphrodites, and that a female could produce *k*_1_ seeds, a male *k*_2_ pollen grains, and a hermaphrodite *fk*_1_ seeds and *mk*_2_ pollen grains. To give these terms some definition, Cruden (2000) reports that pollen-ovule ratios in a sample of animal-pollinated, dioecious, angiosperm species varied from about 10^3^ to 10^6^ with a median of 10,665. While not all ovules necessarily become seeds, it would be reasonable to consider the value of *k*_2_ to be a few orders of magnitude greater than *k*_1_. If the abortion of immature seeds occurs randomly with respect to the sex allocation strategy of the parents, the numbers of pollen and ovules will determine the male and female fitness outcomes.

Charnov (1982) assumed that a hermaphrodite has the same resource availability as a male or female, so that if its entire resource pool were used to make pollen (that is, *f* = 0), it would produce *k*_2_ pollen grains, and similarly *f* = 1 would allow the production of *k*_1_ ovules. To produce both pollen and seeds, *m* and *f* must each be less than unity but their sum may exceed unity. A simple biological circumstance that would allow *m* + *f* > 1 is the “sharing” of fixed start-up costs that must be paid before any reproductive success is obtained. In angiosperms, such fixed costs would include the pedicel, receptacle, sepals and petals of an animal-pollinated flower. Once produced, these floral structures can serve both male and female functions.

#### Fitness

In a population of *n*_1_ + *n*_2_ + *n*_3_ individuals with panmictic mating, any single pollen grain has a probability 1/(*n*_1_*k*_2_ + *n*_3_*mk*_2_) of fertilizing an ovule, and any ovule has a probability 1/(*n*_2_*k*_1_ + *n*_3_*f k*_1_) of fertilizing a pollen grain. The number of zygotes produced by the population is designated *K*. The fitnesses *W*_m_, *W*_f_, and *W*_h_ of an individual male, female, and hermaphrodite can then be specified:

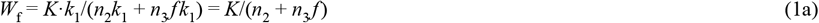

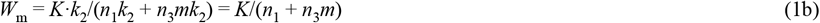

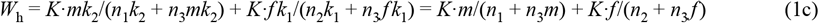

(Charnov 1982, eq. 14.1). The structure of equations (1a-c) deserves some comment. The terms *k*_1_/(*n*_2_*k*_1_ + *n*_3_ *f k*_1_) in equation (1b) and *mk*_2_/(*n*_1_*k*_2_ + *n*_3_*mk*_2_) in equation (1c) represent the number of pollen grains produced by a single male or hermaphrodite, respectively, as a proportion of the total pollen production in the population. Under random panmictic mating, these proportions also indicate the relative reproductive value of the pollen or the “fair share” of total male fitness that an individual will obtain. A corresponding argument applies to ovules and female fitness. *K* haploid genomes are transferred to zygotes via pollen and *K* via ovules, thus defining total fitness for each sex function. *K* can be any positive integer. The only restriction, so obvious that it scarcely attracts notice, is that *K* is a single number, and therefore the same number for male as for female function. When Charnov (1982) transformed *m* and *f* into separate functions representing male and female fitness gain, the required relationship of these functions to *K* was overlooked.

#### Invasion by a hermaphrodite

Charnov (1982) next considers a dioecious population with equal numbers of males and females (*n*_1_ = *n*_2_) and examines the evolutionary conditions under which a rare hermaphrodite can invade. Its fitness would have to exceed that of a male (*W*_h_ > *W*_m_) or a female (*W*_h_ >*W*_f_) or both (although given *n*_1_ = *n*_2_, the average fitness of a male must equal that of a female, and we need consider only one of these inequalities). We can draw on equations (1b, c) to expand *W*_h_ > *W*_m_:

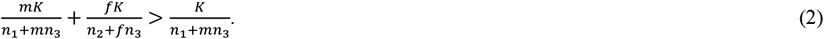

Dividing both sides of the inequality by *K* and ignoring terms with *n*_3_ as being negligible relative to *n*_1_ or *n*_2_, this becomes

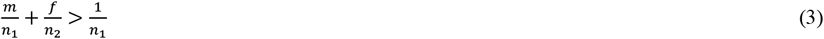

(Charnov 1982, eq. 14.2), and because *n*_1_ = *n*_2_, equation (3) simplifies to *m* + *f* > 1. This is the condition allowing a hermaphrodite to invade a dioecious population. A hermaphrodite must be able to produce more pollen and seed than could be done from the same resource inputs by single-sex plants. As noted already, biologically plausible circumstances, such as sharing an essential fixed start-up cost, would allow this condition to be met. Any values of both *m* and *f* between ½ and 1 satisfy this invasion requirement.

Charnov (1982) then envisions a population of hermaphrodites (*n*_1_ = *n*_2_ = 0), one of which produces 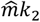 pollen grains and 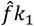 ovules while *n*_3_ common types each produce *mk*_2_ and *f k*_1_. This sets up the standard construction of an evolutionarily stable strategy (ESS) analysis. The fitness of the single mutant can be expressed in a form first used by Shaw and Mohler (1953) in what was recognized later as an early version of an ESS argument (Charnov 1982). I present three slightly different but corresponding versions of the Shaw-Mohler equation:

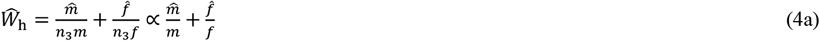

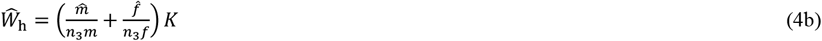

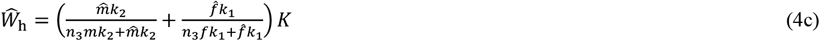

These three forms lead to the same ESS, but they emphasize different aspects of the argument. Equation (4a) is the form presented by Charnov (1982, equation 14.3, p. 221). Strictly speaking, the expression in (4a) calculates the mutant’s relative share of an unspecified total fitness rather than the absolute magnitude of fitness (and also assumes that the contribution of the mutant to the total pool of pollen and ovules is negligible and can be ignored). Equation (4b) simply identifies *K* as the total number of zygotes produced, as in (1), thus making explicit what was unstated in (4a). The specification of *K* emphasizes that the terms *m, f*, 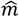 and 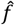 refer to the numbers of pollen and ovules introduced into the mating arena, not the fitness gained from their introduction. Equation (4c) is equivalent to (1c) and includes the pollen and ovule production constants *k*_1_ and *k*_2_ and the contribution of the mutant to the total population pool of pollen and ovules. Constants *k*_1_ and *k*_2_ cancel from the numerators and denominators of the bracketed terms in (4c) but reference to their original use again emphasizes that the terms represent production of pollen and ovules.

#### Gain curves

Charnov (1982, p. 220) asserts that “*m* and *f* actually have more general interpretations[:]…the hermaphrodite has a proportion *f* of a female’s fitness (through seeds) and a proportion *m* of a male’s fitness through pollen.” This seemingly anodyne statement is biologically fraught. It is unobjectionable if it means, for example, that *mk*_2_ pollen grains produced by a hermaphrodite have the same opportunity to mate as any equal number *mk*_2_ out of the *k*_2_ pollen grains produced by a male, and similarly for ovules. That statement is merely an affirmation of “fair competition” for mating between hermaphrodites and single-sex plants. But Charnov’s statement begins to equate *m* and *f* as gamete numbers with a fitness outcome from those gametes, and that is the misstep that leads to error.

Charnov (1982, p. 222) proposes “to plot them (*m, f*) separately, versus a variable like *r* [which represents a resource that can be allocated to male versus female function]. In such a plot *m*(*r*) may be termed the *male gain curve, f* (*r*) the *female gain curve*.” Relying on Bateman’s (1948) principle, Charnov (1982) proposed that *m*(*r*) = *r*^*b*^ and *f* (*r*) = 1 – *r*, in which *r* is the fraction of the reproductive resource pool given to male function. (Charnov denoted the exponent as *n* rather than *b*, but *b* is more clearly distinct from the numbers *n*_1_, *n*_2_, and *n*_3_ that are also part of the argument.) The power function *r*^*b*^ provides a variety of possible shapes, and in particular, diminishing marginal returns when *b* < 1 (Fig. 1). Moreover, these gain curves fulfill the invasion requirement *m* + *f* > 1 for hermaphrodite fitness (equation 3) when *b* < 1; that is, *r*^*b*^ + 1 – *r* > 1. This formulation leads to an ESS at *r*\* = *b*/(1 + *b*) (Charnov 1982).

**Figure 1.**
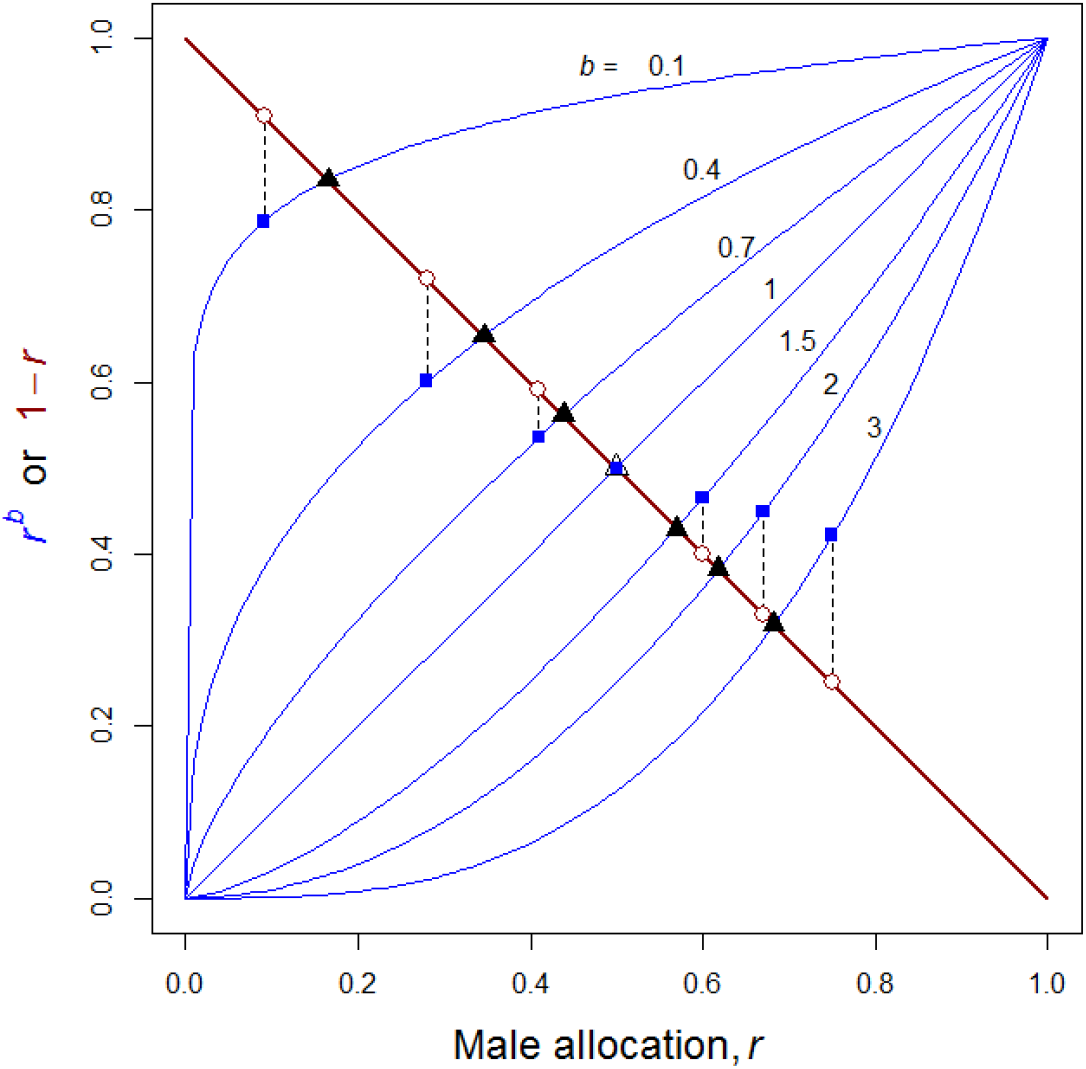
Fitness gain curves and sex allocation optima and in the model of Charnov (1982). Male gain curves, *m* = *r*^*b*^, are shown for several values of *b* as indicated (thin blue curves); the female gain, *f* = 1 – *r*, is shown as a heavy red line. The value of (*r*\*)^*b*^ at the ESS *r*\* = *b*/(1 + *b*) is shown as a filled square; the corresponding value 1 – *r*\* as an open circle. The gap between them is highlighted by a dashed vertical line. The point 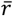 at which *r*^*b*^ = 1 – *r* is shown for each male gain curve as a black triangle. Adapted from Charnov (1982), Figure 14.3.

We must now consider what *m* = *r*^*b*^ and *f* = 1 – *r* mean in the context of this result. If they truly represent male and female fitness outcomes, we can return to equations (4a-c) to remind ourselves that fitness is counted by the number of zygotes of the new generation. Thus, in a population at the ESS equilibrium 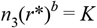 haploid genomes are transmitted via pollen and *n*_3_(1 – *r*\*) = *K* via ovules, which further implies that *r*^*b*^ = 1 – *r*. The solution to this equation has no closed-form expression, but it corresponds to the point of intersection between the *r*^*b*^ and 1 – *r* curves (Fig. 1) and can be calculated numerically. Let us refer to this point as 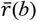 for a given value of *b*. The problem is that 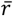 can equal *r*\* = *b*/(1 + *b*) only when *b* = 1; for all other male gain curves, 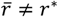 (Fig. 1). If we abandon the putative ESS at *r*\* in favor of 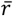 in order to maintain equality of total male and female contributions to the *K* zygotes, fitness is then imposed by fiat rather than derived via fitness maximization under frequency-dependent selection, as the Shaw-Mohler equation and ESS analysis provide.

Alternatively, we can recognize that the fitness interpretation of gain curves was mistaken, despite their label, and allow the functions *m* = *r*^*b*^ and *f* = 1 – *r* to define the production of pollen and ovules. Substitution into the Shaw-Mohler equation in (4c) yields:

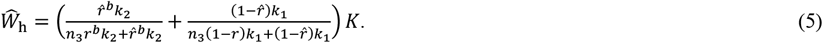

The derivative of (5) evaluated at 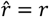 is:

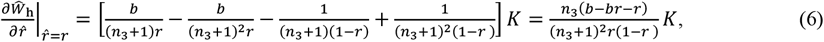

and setting equation (6) equal to zero and solving for *r* yields *r*\* = *b*/(1+*b*). Substitution into equations (4a) or (4b) leads to the same result. Recovery of Charnov’s (1982) ESS from equation (4c) shows that what were called gain curves actually refer to pollen and ovule numbers introduced to the mating arena. Charnov (1982) derived his ESS using a calculation convenience, the MacArthur (1965) product theorem, that further obscured the role of *r*^*b*^ and 1 – *r* as gamete inputs. This is the simple flaw in the gain curve model. What were called fitness gain curves were used in the role of male and female inputs in the framework provided by Shaw and Mohler (1953) and MacArthur (1965).

It is tempting to dismiss this flaw, rebrand gain curves as input curves, and accept the ESS at *r*\* = *b*/(1 + *b*) as legitimate. But then new problems arise. The functional forms *m* = *r*^*b*^ and *f* = 1 – *r* no longer have any justification in Bateman’s principle. Moreover, *r*^*b*^ with *b* < 1 implies that the unit cost of producing and dispersing a pollen grain rises as more and more are produced (or equivalently, that the number of grains produced per unit of resource investment falls as more are produced). It is not clear what biology creates such a circumstance. Moreover, in the absence of Bateman’s principle, there seems no reason that pollen production but not ovule production should be a nonlinear function of resource inputs. Charnov nominated fixed start-up costs of reproduction as a source of the hermaphrodite advantage needed to satisfy the invasion condition in (3), but production of *m* = *r* pollen grains and *f* = (1 – *r*)^*b*^ ovules would also fulfil that condition. And yet under *b* < 1, these new production curves lead to a male-biased ESS allocation while the functions of Charnov (1982) yield a female bias. If optimal sex allocation depends on an arbitrary decision of how a hermaphrodite advantage is distributed between male and female functions, then the generalizations that have emerged from the gain curve theory are suspect.

### Numerical example

A numerical example lacks the generality of an algebraic argument but has the advantage of concrete illustration. Suppose that a parental population comprises *n*_3_ = 10^3^ resident hermaphroditic plants and one mutant. Each plant can produce *k*_2_ = 10^6^ pollen grains if it acts solely as a male or *k*_1_ = 10^3^ ovules if it acts solely as a female (numbers that give a pollen-ovule ratio of the correct order of magnitude for a wide variety of animal-pollinated angiosperms). By acting as a hermaphrodite, however, a resident plant can produce *mk*_2_ = *r*^*b*^(10^6^) pollen grains and *f k*_1_ = (1 – *r*)(10^3^) ovules, and the mutant 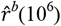 pollen grains and 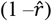 (10^3^) ovules. This set up parallels the development in Charnov (1982, pp. 219-221).

Now set the exponent of the male gain curve at *b* = 0.6. The Charnov ESS would be *r*\* = *b*/(1 + *b*) = 0.375, so let the 1000 resident plants collectively introduce *n*_3_ *mk*_2_ = (10^3^)(0.375^0.6^) (10^6^) = 555,160,759 pollen grains (rounding to the nearest integer) and *n*_3_ *f k*_1_ = (10^3^)(1 – 0.375)(10^3^) = 625,000 ovules to the pollination environment. Inserting these values into Equation (4c) yields

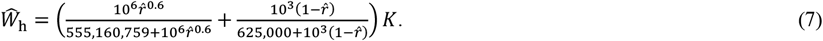

In this example the value of *K* could not exceed the number of ovules introduced to the pollination environment (or, should a different example happen to involve fewer pollen grains than ovules, *K* could not exceed the number of pollen grains). The Shaw-Mohler equation simply assumes *K* is some biologically allowable number and is entirely unconcerned about pollen limitation or seed abortion, so *K* could be smaller than the number of ovules. If the mutant adopts the common allocation 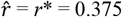, its fitness (to nine decimal places) is given by equation (7) as *Ŵ* _*h*_ = 0.001998002·*K*. A slightly lower male allocation of 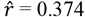 or a slightly higher allocation of 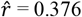 both return a slightly lower mutant fitness of *Ŵ*_*h*_ = 0.001998001·*K*. Plotting *Ŵ*_*h*_ against the entire domain of *r* would revea a single maximum at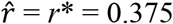. So the Charnov ESS at *r*\* = *b*/(1 + *b*) = 0.375 proves correct in this example, but this is so because equation (7) explicitly used the functions *r*^*b*^ and 1 – *r* to regulate numerical inputs of pollen and ovules to the mating process. Fitness for each sex function at the ESS equilibrium where all plants adopt the same allocation must be *K*, so that mean fitness in a population of 1001 plants could be a maximum of 625 successful pollen grains and 625 mated ovules per plant if there is no pollen limitation, or smaller but equal numbers if not all ovules mate.

### A model with input curves *and* gain curves

It is possible to create a model for hermaphroditic sex allocation in which *r*^*b*^ and 1 – *r* act as pollen and ovule input functions alongside other power functions that genuinely control male and female fitness gains. The basis for such a model is equation (4c), which remains a valid Shaw-Mohler expression; all that is required is an explicit function describing how the outcome of *K* zygotes depends on the true gain curves. To simplify the notation somewhat, I use the labels *P* = *k*_2_ *r*^*b*^ and *O* = *k*_1_ (1 – *r*) for the production of pollen and ovules, respectively, along with their *pro rata* costs of pollen dispersal, pollen receipt and eventual maturation of ovules into seeds.

To allow true gain curves to affect fitness, suppose that a pollination environment of random panmictic mating allows *n*_3_ hermaphrodites to obtain a total of *K* = γ(*n*_3_*P*)^*β*^(*n*_3_*O*)^*δ*^ zygotes. If *β* = *δ* = 1, this mating expression reduces to a conventional model of mass action governed by a parameter γ (fertilization function *F*_1_ in Lehtonen and Dardare 2019). In a pollination context, mass action has a simple biological interpretation. The total number of possible, unique pollen-ovule combinations is the product of *n*_3_*P* and *n*_3_*O*. A pollen grain can mate with only a single ovule, and vice versa, so not all possible pairs can become zygotes, and many pollen grains and ovule may not mate at all. Thus the prevailing pollination environment allows only a small fraction γ of the potential pairs to emerge as successful fertilizations. (A restriction on the value of γ must be added to ensure that the number of zygotes cannot exceed the number of ovules or of pollen—see the Supplemental Material. However, this mating parameter ends up playing no role in the ESS allocation.) The exponents *β* and *δ* add the features of power-function gain curves to the mass-action mating.

Following standard procedures of an ESS analysis, let a single mutant with sex allocation 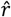 enter the population of *n*_3_ hermaphrodites. It produces 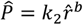 pollen and 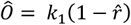 ovules and gains a proportion of the 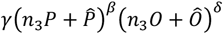 fertilizations that occur based on its relative contribution to the total pool of pollen and ovules in the population. The mutant’s fitness is therefore

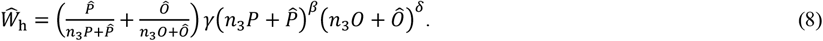

Substituting the explicit expressions for *P*, 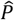, *O*, and 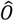 into equation (8), taking the derivative with respect to 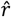, and evaluating the result at 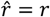 yields

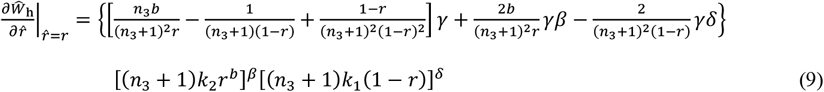

(full derivation in the Supplemental Material). Setting equation (9) equal to zero and solving for *r* yields the ESS candidate

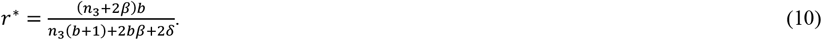

Two additional conditions much be checked to ensure that equation (9) specifies the ESS (Otto and Day 2007); both conditions are satisfied (see the Supplemental Material).

If *β* = *δ* (the gain curves are symmetrical), equation (10) simplifies to *r*\* = *b*/(1 + *b*). Once again, recovery of Charnov’s (1982) result under symmetrical gain curves emphasizes that the original meaning of *m* and *f*—pollen and seed production relative to that of single-sex plants— was correct. Figure 2 represents a population of *n*_3_ = 100 plants, with the pollen production exponent *b* set at 0.9, giving hermaphrodites a realistic advantage over an equivalent single-sex male plant. Under *β* = *δ*, optimal allocation would then be *r*\* ≈ 0.473, indicated by the heavy black line in Fig. 2. Now consider what asymmetrical gain curves (*β* ≠ *δ*) do to sex allocation. The effect is slight, even with considerable disparity between *β* and *δ* (Fig. 2; notice the scale on the vertical axis to appreciate how slight). This trivial effect in Fig. 2 nearly vanishes at *n*_3_ = 1000 and continues to diminish as population size increases (Fig. 3). Given that effective population sizes (*N*_e_) of outcrossing angiosperm species tend to be on the order of 10^3^ to 10^4^ (Schoen and Brown 1991; Gossman et al. 2010), it seems that, as a general rule, gain curves could have no influence on sexual allocation in most populations. Indeed, the small population sizes needed for gain curve asymmetry to have any substantial effect may be too small to persist over sufficient evolutionary time for selection to attain the ESS.

**Figure 2.**
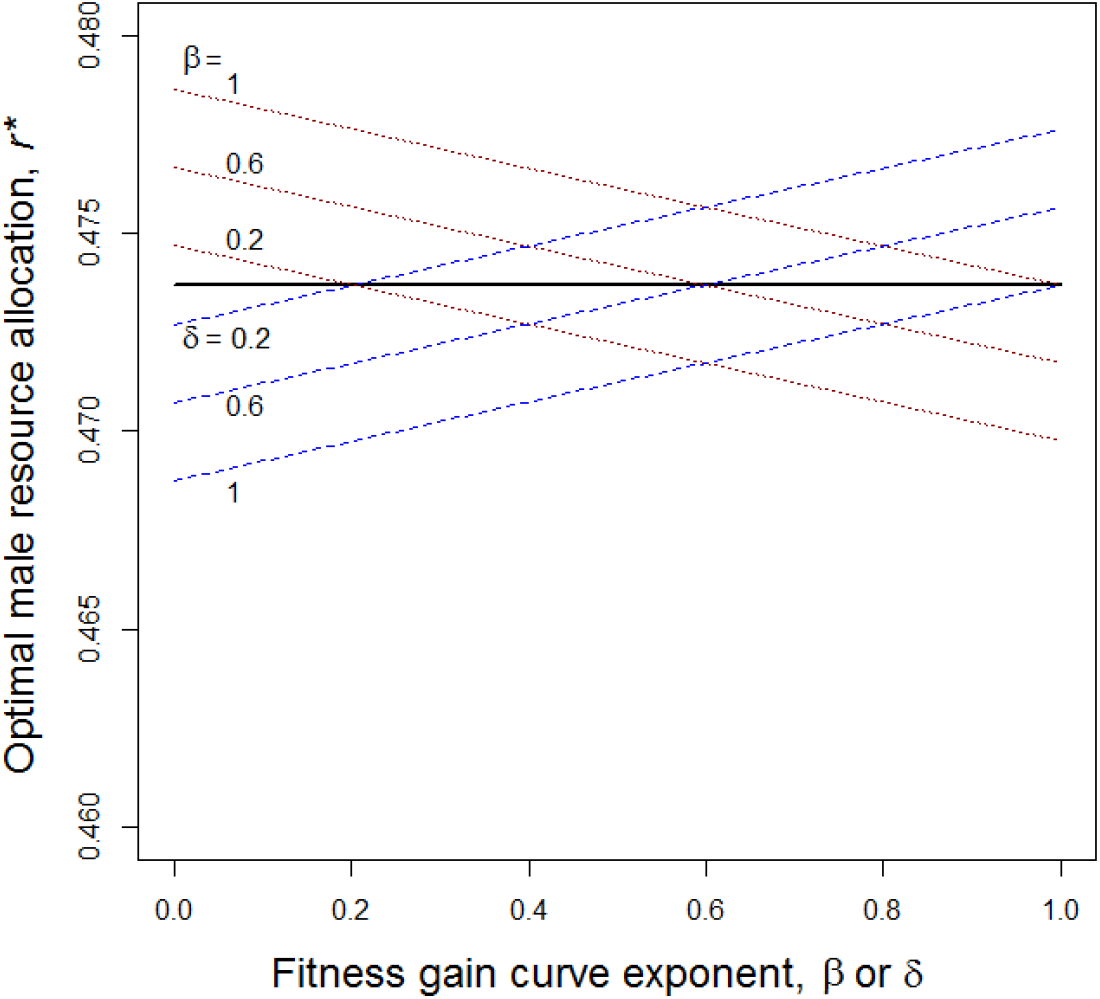
Effect of gain curves on optimal sexual allocation as given by equation (10). Parameters values: *n*_3_ = 100, *b* = 0.9. Thick horizontal black line: *β* = *δ*. Dotted red lines: *β* varies from 0.2 to 1 as indicated in the panel, while *δ* varies as indicated on the horizontal axis. Dashed blue lines: *δ* varies from 0.2 to 1 as indicated, while *β* varies as indicated on the horizontal axis.

**Figure 3.**
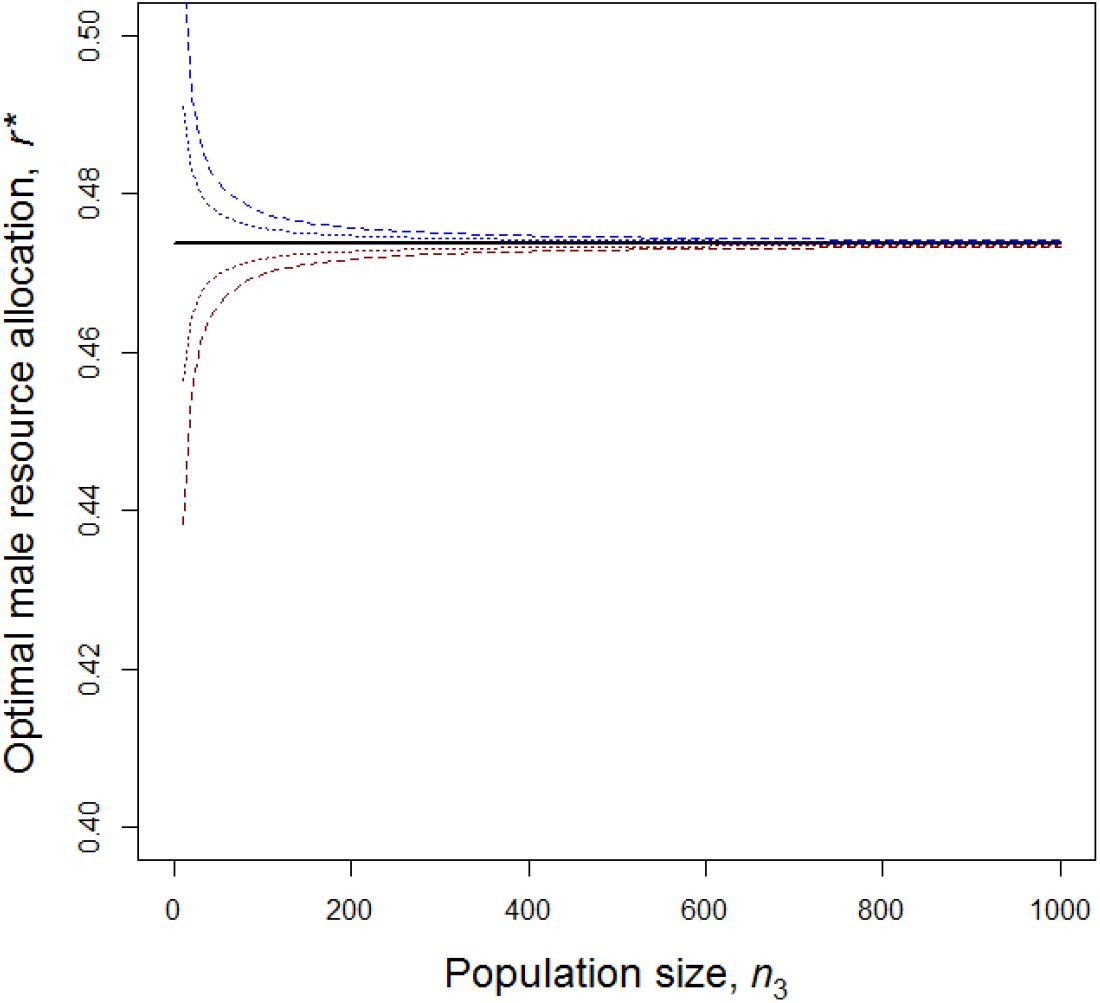
Effect of population size on optimal sexual allocation as given by eqution (10). Parameter for all results: *b* = 0.9. Thick horizontal black line: *β* = *δ*. Dashed blue curve above black line: *β* = 1, *δ* = 0.2. Dotted blue curve above black line: *β* = 1, *δ* = 0.6. Dotted red curve below black line: *β* = 0.6, *δ* = 1. Dashed red curve below black line: *β* = 0.2, *δ* = 1.

## Discussion

The flaw in the original gain curve model was subtle. There was no calculation error. No single defect jumps out of the background. The transition of *m* and *f* from expressions of relative gamete production to fitness gain curves was gradual, with no obviously objectionable step along the way. But although the flaw has a diaphanous character, it is nonetheless serious. We must either abandon the meaning of gain curves as references to fitness outcomes, or repair the original model by separating gamete production from fitness gain, which removes nearly all effect of gain curves.

This will be a difficult realization to digest in a field that has long taken the explanatory power of gain curves for granted. Lack of causal efficacy will be all the more difficult to accept because gain curves as descriptions of reproductive ecology must exist, as noted in the Introduction. But descriptions are not causes, and, in retrospect, the problem with specifying male fitness independently of female fitness is readily perceived. New explanations for many sex allocation patterns that have been attributed to gain curves will be needed.

I will used wind pollination as an example of how such a reassessment of gain curves might proceed. Male-biased allocation has been found in some wind-pollinated angiosperms (Lemen 1980; McKone 1987; McKone et al. 1998) and a pronounced male bias has also been found in the wind-dispersed, heterosporous lycophyte *Selaginella* (Petersen and Burd 2018). It is thought that wind cannot be saturated as a pollen transport vector in the way that animal vectors can, and so the male bias is commonly attributed to linear or nearly linear male fitness gains under wind dispersal (Charnov 1979; Charlesworth and Charlesworth 1981; Friedman and Barrett 2011; Aljiboury and Friedman 2022). Release of pollen higher above the ground should further enhance the efficacy of wind as a pollen vector, and accordingly male flower production has been found to increase with plant height in several diclinous wind-pollinated species (McKone and Tonkyn 1986; Burd and Allen 1988; Solomon 1989; Bickel and Freeman 1993; Fox 1993; Friedman and Barrett 2011). Male-biased allocation in wind-pollinated plants corresponds intuitively and appealingly to the gain-curve explanation. Yet equation (10) implies that this explanation cannot be correct.

What, then, does cause male-biased allocation under wind dispersal? A likely explanation (Petersen and Burd 2018) lies in sexual differences in dispersal. Bulmer and Taylor (1980) suggested that such differences should have a general effect on sex allocation, favoring allocation to the sex function that enjoys better dispersal. Fromhage and Kokko (2010) have since confirmed this suggestion in a more thorough analysis. The effect of dispersal arises because a larger dispersal kernel decreases the degree of competition among the pollen or seeds from a single plant and simultaneously increases their competition with non-relatives. Fromhage and Kokko (2010) showed that the potential effect on sex allocation is quite large, leading to very highly male-biased or highly female-biased allocation depending on the dispersal circumstances. In contrast, the model leading to equation (10) above specifically avoids all dispersal effects with its mass-action representation of panmictic mating, and optimal sexual allocation accordingly remains near equality (Figs. 2 and 3), deviating only under the effect of the production parameter *b*. Since spatial structuring of fitness opportunities based on dispersal seems likely to be inevitable in most plant populations, the argument of Fromhage and Kokko (2010) may provide a general explanation for broad patterns of sexual resource allocation that deviate from male-female equality.

Thus, male-biased allocation under wind-dispersal can be explained if pollen or microspores are usually better dispersed than seeds or megaspores. This seems to be the case. Wind can transport viable pollen over tens and hundreds of kilometers (Williams, 2010; Buschbom et al., 2011; Kremer et al., 2012) and it has been recognized that the inefficiency commonly attributed to wind pollination may be mistaken (Friedman and Barrett 2009). Linear gain curves may describe the nearly inexhaustible capacity of wind to carry pollen, but sexual dispersal differences (and the ensuing differences in relative intra- and inter-cohort competition) might explain the resulting sexual allocation.

## Conclusion

Should we despair that gain curves lack the explanatory power we had supposed? I think not—indeed, the dispersal effects identified by Bulmer and Taylor (1980) and Fromhage and Kokko (2010) raise new, unexplored possibilities for investigation. Some very large issues might profitably be addressed. Angiosperm reproduction is noteworthy for its delayed investment in endosperm and fruit tissue until after ovule fertilization has occurred (Floyd and Friedman 2000). This reproductive pattern leads to a pronounced female bias in end-of-season sex allocation in most species (Cruden and Lyon 1985; Goldman and Willson 1986; Campbell 2000). The contribution of endosperm to seedling success is well recognized (Baroux et al. 2002; Moles and Westoby 2004; Lafon-Placette and Köhler 2014) and the protective and dispersal advantages of carpels and fruit tissues have received decades of study (Lorts et al. 2008; Valenta and Nevo 2020; Lei et al. 2021). The ecological advantages of endosperm and fruit may be so evident that their concomitant role in female allocation has raised few questions. Yet these fundamental features of the angiosperm life cycle have never been explained from the perspective of sexual allocation. Are they so advantageous that selection favored their appearance at the origin of the angiosperm lineage (Scutt et al. 2006) despite an otherwise maladaptive sexual allocation that they entailed? Or did (and does) the selective regime favor a female bias, and if so, why?

The ideas of Fromhage and Kokko (2010) offer a potential explanation. Their model treated dispersal in terms of distance but the logic of their argument would accommodate dispersal through time via seed dormancy. If establishment opportunities emerge and vanish dynamically, then time is a dimension of their availability in the environment and seed dormancy is part of dispersal relative to the “density” of fitness opportunities in the environment. Dormancy may give seed dispersal a nearly universal advantage over pollen dispersal and provide a general explanation of the common female-biased sexual allocation in angiosperms I believe that a link between sexual allocation and seed dormancy has never been investigated, but the Fromhage and Kokko (2010) model points in that direction.

The transition away from gain curves to new explanations of sex allocation may be straightforward, as it appears at first glance to be in the case of wind dispersal. Other cases may prove more difficult. But new insights will arise when causal efficacy is correctly placed.

## Acknowledgements

I thank Eric Charnov for generously providing comments on these ideas.

