## Supplemental Material for "The concept of the gain curve"

Draft: 7 January 2024

### Supplemental Material

This supplement provide a detailed derivation of the evolutionarily stable strategy (ESS) in the model outlined in the main text. A candidate for the ESS sex allocation, *r**, must satisfy three conditions (Otto and Day 2007, Equations 12.15b, c, d):

1. The first derivative of the mutant's fitness function with respect to $\hat{r}$, evaluated at $\hat{r} = r$, must equal zero; that is, $\left.\frac{\partial \widehat{W}_{\mathrm{h}}}{\partial \hat{r}}\right|_{\hat{r}=r} = 0$. This is the usual necessary albeit insufficient condition for the maximum of a function.

2. The fitness gradient given by $\partial \widehat{W}_{\mathrm{h}}/\partial \hat{r}\big|_{\hat{r}=r}$ must itself have a negative derivative at *r**, $\frac{d}{dr}\left(\left.\frac{\partial \widehat{W}_{\mathrm{h}}}{\partial \hat{r}}\right|_{\hat{r}=r}\right)_{r=r^*} < 0$. This condition, called convergence stability, ensures that selection will favor greater male allocation if the common allocation in the population is below the optimum, *r*

< $r$*, and reduced male allocation if the common alloction exceeds the optimum, $r > r$*, but cease to act at $r = r$*.

3. The second derviative, evaluated at $\hat{r} = r = r^*$, must be less than or equal to zero, $\left.\frac{\partial^2 \widehat{W}_{\mathrm{h}}}{\partial \hat{r}^2}\right|_{\hat{r}=r=r^*} \leq 0$. This condition insures that $r$* corresponds to a maximum in the fitness function rather than a minimum or inflection point.

To ensure that the mating function of equation (8) does not yield more zygotes than the number of ovules available, we must restrict the parameter $\gamma$ so that $\gamma\left(n_3 P + \hat{P}\right)\left(n_3 O + \hat{O}\right) < n_3 O + \hat{O}$, or $\gamma < 1/\left(n_3 P + \hat{P}\right)$. A similar restriction $\gamma < 1/\left(n_3 O + \hat{O}\right)$ is also required to ensure that zygote formation does not exceed the number of pollen grains, but normally the ovule limit will be more restrictive.

Equation (8) of the main text presents the mutant fitness function:

$$\widehat{W}_{\mathrm{h}} = \left(\frac{\hat{P}}{n_3 P + \hat{P}} + \frac{\hat{O}}{n_3 O + \hat{O}}\right)\gamma\left(n_3 P + \hat{P}\right)^{\beta}\left(n_3 O + \hat{O}\right)^{\delta}. \qquad (8)$$

$P$ and $\hat{P}$ are labels for $k_2 r^b$ and $k_2 \hat{r}^b$ pollen grains; $O$ and $\hat{O}$ for $k_1(1 - r)$ and $k_1(1 - \hat{r})$ ovules. Making the substitution into equation (8) yields

$$\widehat{W}_{\mathrm{h}} = \left(\frac{k_2 \hat{r}^b}{n_3 k_2 r^b + k_2 \hat{r}^b} + \frac{k_1(1-\hat{r})}{n_3 k_1(1-r) + k_1(1-\hat{r})}\right)\gamma(n_3 k_2 r^b + k_2 \hat{r}^b)^{\beta}\left(n_3 k_1(1-r) + k_1(1-\hat{r})\right)^{\delta}, \quad \text{(S1)}$$

and with some simplifiction this becomes

$$\widehat{W}_{\mathrm{h}} = \left(\frac{\hat{r}^b}{n_3 r^b + \hat{r}^b} + \frac{1-\hat{r}}{n_3(1-r)+1-\hat{r}}\right)\gamma k_1 k_2 (n_3 r^b + \hat{r}^b)^\beta (n_3(1-r)+1-\hat{r})^\delta. \tag{S2}$$

Finding the derivative of (S2) "by hand" is straightforward but tedious. It can also be determined with machine assitance from programs such as *Mathcad* or *Mathematica* or similar. I used *Mathcad*. With some minor simplification the derivative is:

$$\frac{\partial \widehat{W}_{\mathrm{h}}}{\partial \hat{r}} =$$

$$\left\{\left[\frac{b\hat{r}^{b-1}}{n_3 r^b + \hat{r}^b} - \frac{b}{\hat{r}}\left(\frac{\hat{r}^b}{n_3 r^b + \hat{r}^b}\right)^2 - \frac{1}{n_3(1-r)+1-\hat{r}} + \frac{1-\hat{r}}{[n_3(1-r)+1-\hat{r}]^2}\right] + \right.$$

$$\left.\left[\beta\frac{b\hat{r}^{b-1}}{n_3 r^b + \hat{r}^b} - \delta\frac{1}{n_3(1-r)+1-\hat{r}}\right]\left[\frac{\hat{r}^b}{n_3 r^b + \hat{r}^b} + \frac{1-\hat{r}}{n_3(1-r)+1-\hat{r}}\right]\right\}\gamma k_1 k_2 (n_3 r^b + \hat{r}^b)^\beta [n_3(1-r)+1-\hat{r}]^\delta\ .$$

(S3)

Setting $\hat{r} = r$ with some further minor simplification yields equation (8) of the main text:

$$\left.\frac{\partial \widehat{W}_{\mathrm{h}}}{\partial \hat{r}}\right|_{\hat{r}=r} = \left\{\left[\frac{n_3 b}{(n_3+1)^2 r} - \frac{1}{(n_3+1)(1-r)} + \frac{1-r}{(n_3+1)^2(1-r)^2}\right]\gamma + \frac{2b}{(n_3+1)^2 r}\gamma\beta - \frac{2}{(n_3+1)^2(1-r)}\gamma\delta\right\}\cdot$$

$$[(n_3+1)k_2 r^b]^\beta [(n_3+1)k_1(1-r)]^\delta. \tag{8}$$

Now consider the ESS conditions:

1. *First derivative condition*

Equation (8) will equal zero if

$$\left[\frac{n_3 b}{(n_3+1)^2 r} - \frac{1}{(n_3+1)(1-r)} + \frac{1-r}{(n_3+1)^2(1-r)^2}\right]\gamma + \frac{2b}{(n_3+1)^2 r}\gamma\beta = \frac{2}{(n_3+1)^2(1-r)}\gamma\delta. \qquad \text{(S4)}$$

Eliminating γ from both sides of the equality in (S4), followed by some simplification of terms yields

$$\frac{(n_3+2\beta)b}{(n_3+1)^2 r} - \frac{1}{(n_3+1)(1-r)} + \frac{1-r}{(n_3+1)^2(1-r)^2} + \frac{2b\beta}{(n_3+1)^2 r} = \frac{2\delta}{(n_3+1)^2(1-r)}. \qquad \text{(S5)}$$

Multiplying both sides of the equality in (S4) by $(n_3 + 1)^2(1 - r)/(2\delta)$ yields

$$\frac{(n_3+2\beta)b}{(n_3+1)^2 r} \cdot \frac{1-r}{\delta r} - \frac{n_3+1}{2\delta} + \frac{1}{2\delta} = 1, \qquad \text{(S6)}$$

$$\frac{(n_3+2\beta)b}{2} \cdot \frac{1-r}{\delta r} = 1 + \frac{n_3}{2\delta}, \qquad \text{(S7)}$$

$$\frac{1-r}{r} = \left(1 + \frac{n_3}{2\delta}\right)\frac{2\delta}{(n_3+2\beta)b}, \qquad \text{(S8)}$$

leading finally to equation (10) of the main text:

$$r^* = \frac{(n_3+2\beta)b}{n_3(b+1)+2b\beta+2\delta}. \qquad (10)$$

2. *Convergence stability*

The derivative of the fitness gradient is

$$\frac{d}{dr}\left(\frac{\partial \widehat{W}_{\mathrm{h}}}{\partial \hat{r}}\bigg|_{\hat{r}=r}\right) =$$

$$\left\{\left[-\frac{n_3 b}{(n_3+1)^2 r^2}-\frac{n_3+2}{[n_3(1-r)+1-r]^2}+\frac{2(n_3+1)(1-r)}{[n_3(1-r)+1-r]^3}\right]\gamma-\frac{2b}{(n_3+1)^2 r^2}\beta\gamma-\frac{2}{(n_3+1)^2(1-r)^2}\delta\gamma+\right.$$

$$\left.\left[\left(\frac{n_3 b}{(n_3+1)^2 r}-\frac{1}{n_3(1-r)+1-r}+\frac{1-r}{[n_3(1-r)+1-r]^2}\right)\gamma+\frac{2b}{(n_3+1)^2 r}\beta\gamma-\frac{2}{(n_3+1)^2(1-r)}\delta\gamma\right]\left(\frac{b\beta}{r}-\frac{\delta}{1-r}\right)\right\}[(n_3+$$

$$1)k_2 r^b]^\beta[(n_3+1)k_1(1-r)]^\delta. \qquad \text{(S9)}$$

Numerical evaluation suggests that the expression in (S9) is everywhere negative for biologically meaningful values of the parameters, but for convergence stability we need only evaluate this derivative at the candidate ESS, $r^* = (n_3 + 2\beta)b/[n_3(b + 1) + 2\beta b + 2\delta]$. Substituting this for $r$ in (S9) and simplifying yields

$$\frac{d}{dr}\left(\frac{\partial \widehat{W}_{\mathrm{h}}}{\partial \hat{r}}\bigg|_{\hat{r}=r}\right)_{r=r^*} =$$

$$-\frac{[n_3(b+1)+2b\beta+2\delta]^3\gamma}{(n_3+2\delta)(n_3+1)^2(n_3+2\beta)b}\cdot\left[(n_3+1)k_2\left(\frac{(n_3+2\beta)b}{n_3(b+1)+2b\beta+2\delta}\right)^b\right]^\beta\cdot\left[\frac{(n_3+1)k_1(n_3+2\delta)}{n_3(b+1)+2b\beta+2\delta}\right]^\delta. \qquad \text{(S10)}$$

Since all the parameters in (S10) have only positive values (to be biologically meaningful), the expression as a whole is necessarily negative, and convergence stability is satisfied.

3. *Second derivative condition*

The second derivative of equation (S2) is a lengthy expression. We need consider its value only at the candidate ESS, $\hat{r} = r = r^*$, which provides a degree of simplification. Substituting the value $r^* = (n_3 b + 2b\beta)/[n_3(b+1) + 2b\beta + 2\delta]$ into $\partial^2 \widehat{W}_{\mathrm{h}}/\partial \hat{r}^2$ and simplifying yields an expression of the form

$$\left.\frac{\partial^2 \widehat{W}_{\mathrm{h}}}{\partial \hat{r}^2}\right|_{\hat{r}=r=r^*} = (A + \mathrm{B} + \mathrm{C} + \mathrm{D}) \cdot \mathrm{E} \tag{S11}$$

in which

$$\mathrm{A} = \frac{n_3(b-1)b-1}{(n_3+1)(n_3+2\beta)^2} \cdot \frac{n_3[n_3(b+1)+2b\beta+2\delta]^2}{b} - \frac{2n_3}{(n_3+1)^3} \cdot \frac{[\,n_3(b+1) + 2b\beta + 2\delta]^2}{(n_3+2\delta)^2} \tag{S11a}$$

$$\mathrm{B} = \left[\frac{b-1}{b} \cdot \frac{\beta[n_3(b+1)+2b\beta+2\delta]^2}{(n_3+2\beta)^2}\left(\frac{1}{n_3+1} - \frac{1}{(n_3+1)^2}\right) - \frac{\delta}{(n_3+1)^2} \cdot \frac{[\,n_3(b+1) + 2b\beta + 2\delta]^2}{(n_3+2\delta)^2}\right]\frac{2}{n_3+1} \tag{S11b}$$

$$\mathrm{C} = -\frac{2n_3^2[\,n_3(b+1) + 2b\beta + 2\delta]^2(\beta-\delta)^2}{(n_3+1)^3(n_3+2\beta)^2(n_3+2\delta)^2} \tag{S11c}$$

$$\mathrm{D} = \left[-\frac{2n_3(\beta-\delta)[n_3(b+1) + 2b\beta + 2\delta]}{(n_3+1)^2(n_3+2\beta)(n_3+2\delta)} - \frac{n_3(\delta-\beta)[n_3(b+1) + 2b\beta + 2\delta]}{(n_3+1)(n_3+2\beta)(n_3+2\delta)}\left(1 + \frac{1}{n_3+1} + \frac{(n_3+2\delta)^2}{n_3[\,n_3(b+1) + 2b\beta + 2\delta]^2} - \frac{(n_3+2\beta)b}{n_3(b+1) + 2b\beta + 2\delta}\right)\right]\frac{\beta}{n_3+1} \cdot \frac{n_3(b+1) + 2b\beta + 2\delta}{n_3+2\beta} \tag{S11d}$$

$$E = k_1 k_2 \gamma \left[(n_3+1)\left(\frac{(n_3+2\beta)b}{n_3(b+1) + 2b\beta + 2\delta}\right)^b\right]^\beta \left[(n_3+1)\left(1 - \frac{(n_3+2\beta)b}{n_3(b+1) + 2b\beta + 2\delta}\right)\right]^\delta. \tag{S11e}$$

The second derivative condition will be satisfied because (A + B + C +D) is always negative and E is always positive, leaving the overall expression (S11) negative. We need consider only situations in which $b < 1$, since $b > 1$ leaves hermaphrodites at a distadvantage in gamete production relative to males and females, so that hermaphroditism would not be stable.

The sign of A is determined by the term ($b – 1$), which will be zero when $b = 1$ and negative for $b < 1$, leaving the entire expression in (S11a) negative for $b < 1$. Note also that the absolute value of the first term on the right-hand side of the equality will be on the order of magnitude of unity, because $n_3$ appears as a factor the same number of times in the numerator and denominator. The absolute value of the second term on the right-hand side, in contrast, will be much smaller, on the order of $n_3^3/n_3^5 = n_3^{-2}$. The absolute value of A governs the value of (A + B + C +D) as a whole.

The value of B will similarly be negative for $b < 1$ because of the term ($b – 1$) in (S11b), but its absolute value will be small relative to A.

The parameter values making up (S11c) are all positive, to be biologically meaningful, and so the overall value of C is inherently negative.

D equals zero when $\beta = \delta$ because of the terms ($\beta – \delta$) and ($\delta – \beta$) in (S11d). When the gain curve exponents are unequal, D can take on positive or negative values, depending on the nature of the gain curve asymmetry. But even when positive, the absolute magnitude of D will be small relative to A, because $n_3$ appears as a factor more often in the denominator than in the numerator of the ratios appearing in (S11d). Given that A, B and C all take negative values and that D will have small absolute value whether positive or negative, the sum (A + B + C + D) is always negative.

In (S11e), the ratio $(n_3 b + 2b\beta)/[n_3(b+1) + 2b\beta + 2\delta]$ (i.e, the value of $r^*$) will always be less than unity, so that $(1 - (n_3 b + 2b\beta)/[n_3(b+1) + 2b\beta + 2\delta])$ will always be positive. Then given positive values for all parameters, expression $E$ will always be positive, leaving the entire second derivative in (S11) always negative. The second-order condition is therefore satisfied, and $r^*$ represents a fitness maximum.
